# Habitual ground nesting in the Bugoma Forest chimpanzees (*Pan troglodytes schweinfurthii*), Uganda

**DOI:** 10.1101/2022.12.14.520400

**Authors:** Catherine Hobaiter, Harmonie Klein, Thibaud Gruber

## Abstract

We report the presence of habitual ground nesting in a newly studied East African chimpanzee *(Pan troglodytes schweinfurthii)* population in the Bugoma Central Forest Reserve, Uganda. Across a 2-year period we encountered 891 night-nests, 189 of which were classified as ground nests, a rate of ∼21%. We find no preliminary evidence of socio-ecological factors that would promote its use and highlight local factors, such as high incidence of forest disturbance due to poaching and logging, which appear to make its use disadvantageous. While further study is required to establish whether this behaviour meets the strict criteria for non-human animal culture, we support the argument that the wider use of population and group-specific behavioural repertoires in flagship species, such as chimpanzees, offers a tool to promote the urgent conservation action needed to protect threatened ecosystems, including the Bugoma forest.

## Introduction

All apes construct overnight arboreal nests by interweaving branches and other vegetation (Fruth et al., 2018; Anderson et al., 2019). Gorillas, although able to climb while foraging, are the most terrestrial of the great apes and build overnight nests primarily on the ground using available herbaceous vegetation, although they increase the use of arboreal nesting seasonally, or when terrestrial vegetation is limited (Tutin et al., 1995; Brugiere & Sakom, 2000; Mfossa et al., 2022). In contrast, orangutans, bonobos, and chimpanzees primarily build arboreal nests (Goodall 1962; Fruth & Hohmann, 1993; Prasetyo et al., 2009; Hicks, 2010; van Casteren et al., 2012; Fruth et al., 2018), using multiple interwoven supporting branches, filling in with smaller softer material (Furuichi & Hashimoto, 2000; Prasetyo et al., 2009; Samson & Hunt, 2012; Fruth et al., 2018). There may be an effect of learning, with younger apes constructing more robust nests earlier if exposed to adults who build nests (orangutans: Lethmate, 1977; chimpanzees: McGrew, 2004; Videan, 2006), but further work with wild populations is needed to more carefully disentangle potential social learning from ecological and developmental influences. Each independent individual constructs their own nest, but individuals will typically nest in a party with others in the same or near-by trees (Fruth, 1995; Schaller, 1963; Goodall, 1968). Chimpanzees and bonobos typically construct nests at between 8 to 20m in height, but they may be as low as 3-4m typically where taller trees are limited (Pruetz et al., 2008; Samson & Hunt, 2012; Fruth et al., 2018). Full nests – night or day – are distinguished in their construction from other flimsier resting structures such as ‘day beds’ and ‘cushions’, which typically involve only 1-2 bent branches, loosely interwoven, and may be as simple as a single small sapling bent over or a clump of ferns (Boesch, 1995; Furuichi & Hashimoto, 2000; Brownlow et al., 2001; Koops et al., 2007). Nests are constructed each evening for overnight use, but they are also built during the day for a range of reasons from sleeping, to play, or sexual solicitation (Boesch, 1995; Plumptre and Reynolds, 1997; Fruth and Hohmann, 1996; Brownlow et al., 2001; McGrew, 2010; Fruth et al., 2018), with bonobos also regularly constructing additional full nests during the day (Wessling & Surbeck 2022).

Despite the use of arboreal night nesting in every chimpanzee community studied to date, our understanding of their function remains limited, with – potentially complementary – hypotheses currently including seasonal patterns (Samson & Hunt, 2014; van Casteren et al., 2012), thermoregulation (Koops et al., 2012; McGrew, 2004; Stewart et al., 2018), pathogen and parasite avoidance (Anderson, 1998; Lacroux et al., 2022), and predation avoidance (Koops et al., 2012; Kortlandt, 1992; Pruetz et al., 2008; Stewart & Pruetz, 2013). Some aspects of the local environment appear to impact the choice of nest location, for example a preference for slopes over flatter ground (Issa: Hernandez-Aguilar 2009; Mahale: Izawa and Itani, 1966; Suzuki, 1969; Assirik: Baldwin, 1979). However, in habitats where trees are limited this tends to lead to a concentration of nests in available trees (Hernandez-Aguilar 2009), rather than alternative strategies such as ground nesting.

The very occasional use of overnight ground nests is widely reported in all chimpanzee sub-species, including in Uganda (see below and c.f. Tagg et al., 2013). Excluding the minimally constructed day beds and cushions (Furuichi & Hashimoto, 2000; Koops et al., 2007), the use of fully constructed overnight nests on the ground is typically rare (5 to 10% of nests constructed in a given population; Matsuzawa & Yamakoshi, 1996; Koops et al. 2007), perhaps as a fall-back by sick or injured individuals (<1%; Furuichi & Hashimoto, 2000). Slightly more frequent use of overnight ground nests was reported by male West African chimpanzees (*Pan troglodytes verus*) of Seringbara in the Nimba Mountains in Guinea (∼3% to 5% of night nests; Koops et al., 2007; 2012) and of Fongoli long term field site (∼12% of night nests; Stewart 2011). Frequent ground nests have also been recently observed in Cameroun in Nigeria-Cameroun chimpanzees (*Pan troglodytes ellioti*) of one (i.e., Andu) of the two field sites of the Lebialen-Mone Forest (∼32% of night nests; Last & Muh, 2013) and in Central African chimpanzees (*Pan troglodytes troglodytes)* of La Belgique research field site in the Dja Biosphere Reserve (∼3% to ∼9% of night nests; Guislain & Duplain, 2005; Tagg et al., 2013). However, despite ∼ 250 years of continuous observations across long-term research sites (Gombe, >60 years; Mahale, >60 years; Kibale-Kanyawara, >30 years; Kibale-Ngogo, >20 years; Kalinzu, >30 years; Budongo-Sonso, >30 years, Budongo-Waibira, >10 years); the customary use of overnight ground nests has only been reported in East African chimpanzee communities in the Northern Democratic Republic of Congo (∼11% of nesting sites; Hicks et al., 2019).

Here we report on the presence of apparently habitual ground nesting in the newly studied East African chimpanzee *(Pan troglodytes schweinfurthii)* population in the Bugoma Central Forest Reserve, in Uganda. In addition, while we consider the specific socio-ecological factors that would promote its use, we suggest that local factors such as extended anthropogenic disturbance would be more likely to inhibit the building of ground nests, and highlight the possibility of a cultural explanation. Establishing a case for cultural behaviour on the basis of either evidence of social transmission or the exclusion of alternative hypotheses, such as genetic or ecological variation may take decades in long-lived species. However, we argue that the use of group-differences in the behavioural repertoires in flagship species, including chimpanzees – irrespective of the socio-ecological or cognitive mechanism by which they are produced – offers an effective tool to promote the urgent conservation action needed to protect threatened ecosystems, such as the Bugoma forest.

## Methods

### Site

The Bugoma Central Forest Reserve includes 400km^2^ of semi-deciduous tropical rain forest (located at 01°15′N 30°58′E and at 990-1300m elevation). Located between the Central Forest Reserve of Budongo (425km^2^) and Kibale National Park (776km^2^), Bugoma represents the largest contiguous forest habitat for chimpanzees in East Africa in which there has been no long-term research activity. In 2006, chimpanzee density was estimated at 1.9 chimpanzees per km^2^ (giving an approximate population estimate of ∼760 chimpanzees; Plumptre and Cox, 2006) but given widespread primate population declines (Estrada et al., 2017) a current population estimate of ∼600 chimpanzees may be more appropriate. Following surveys in 2015, in 2016 the Bugoma Primate Conservation Project (BPCP, www.bugomaprimates.com) initiated daily habituation and monitoring activities across several chimpanzee communities. Our data are concentrated on the Mwera community, where regular contacts were initiated in 2016 and were typically occurring on most days by 2018. Chimpanzees in the Mwera-North community were also encountered, as their range overlaps with that of Mwera, but they are not intentionally followed and they still tend to flee when our presence is detected allowing us to distinguish between the groups. The current estimated range for the Mwera community is ∼10km2 (see Figure 1). During the study period, chimpanzees in the Mwera community were followed on regular basis (several days a week for several hours per day) with an estimate of 60-80 individuals based on local East African chimpanzee community sizes (Wilson et al., 2014). However, as it was not possible to nest or denest them, direct observations of night nest building were not available.

**Figure 1:**
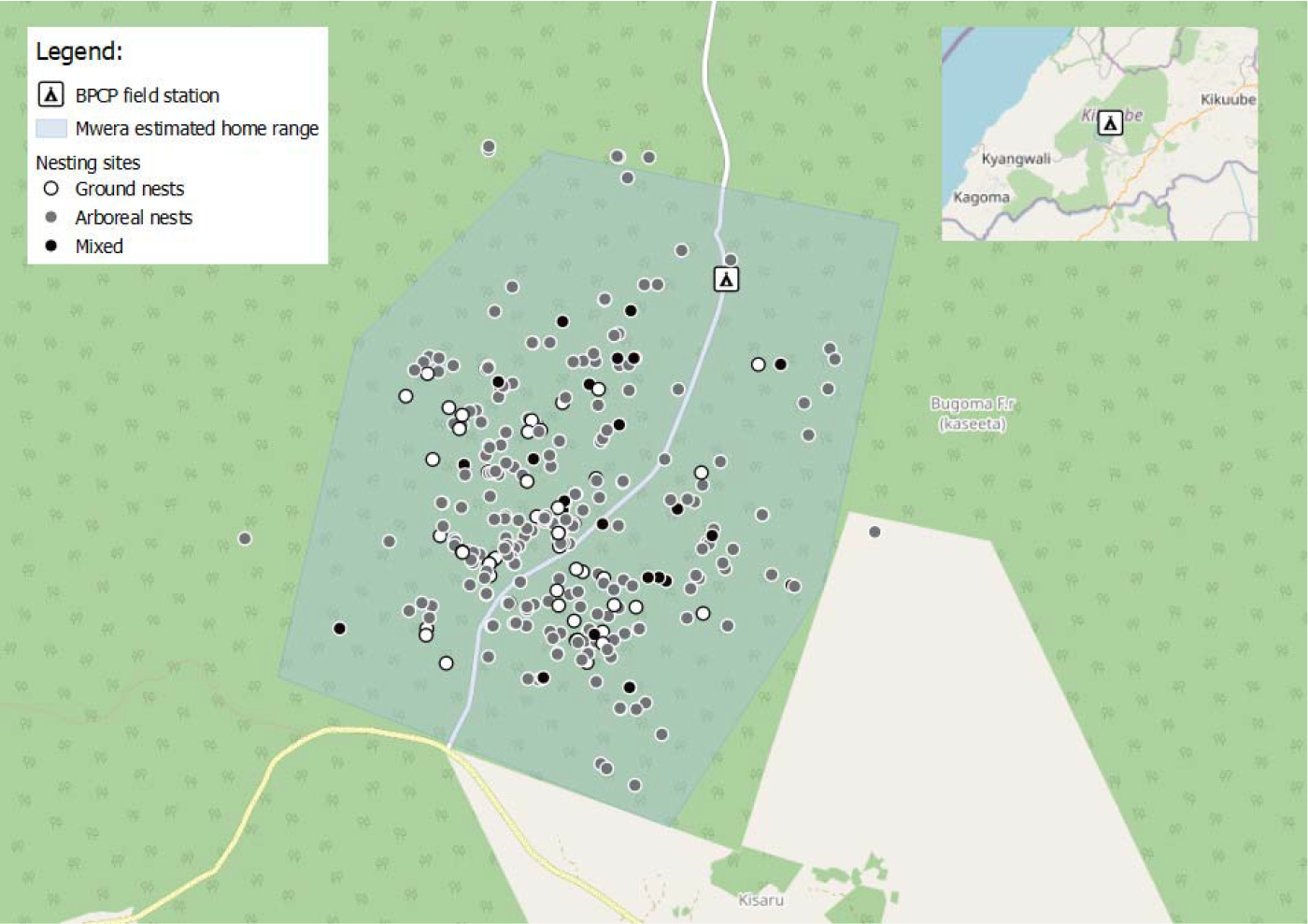
Location of nesting sites within the current estimated range of the Mwera chimpanzee community in Bugoma Central Forest Reserve. White = ground nest only site; Grey = tree nest only site; Black = ground and tree nests in same site. Inset shows full boundaries of Forest Reserve.

### Data collection

In addition to direct observations of chimpanzee behavioural and health data, project staff record the type and GPS location of signs of chimpanzee activity (prints, dung, feeding remains, discarded tools, and nests), and of illegal human activity (primarily hunting and logging) whenever they are encountered ad libitum.

A ‘nesting site’ was defined as one or more nests of the same age in which no two nests were more than 30m apart (Furuichi & Hashimoto, 2000). Following increasing encounters with overnight ground nests, in 2019, BPCP staff started to collect detailed information on overnight nests. However, data collection was impacted by the Covid19 outbreak with 1) a stop of data collection from February 2020 to July 2020 and 2) a change of data collection protocol after this period. Therefore, here, we provide the following nesting data: GPS location, approximate age (1 day, 2 days, under a week, over a week), the nest type (ground, tree) and the number of ground and tree nests in the nesting site from August 2019 to June 2022. Nest decay rates vary substantially between populations and are shaped by local ecological and seasonal factors (Morgan et al., 2016; Wessling & Surbeck, 2021). We followed Romani and colleagues (2023); who recently described nest stages and decay rates for the Bugoma Central Forest Reserve. Other nesting data, including: density of surrounding forest (Open, Medium, Dense), forest type (primary, secondary-mixed, disturbed, water-swamp, grassland), the height from the ground of all tree nests (m), the primary tree species used, the width and length of ground nests (cm), and the use of bent or detached branches and the inclusion of Terrestrial Herbaceous Vegetation (hereafter: terrestrial herbs) in ground nest construction, are only available for the pre-Covid19 outbreak, i.e. from August 2019 to February 2020.

### Data availability

All data used for the analyses and figures in this manuscript are available in a publicly accessible GitHub repository: https://github.com/Wild-Minds/GroundNesting_Bugoma

### Ethical statement

This study was observational and did not involve any interventions, apart from daily visits to the chimpanzee communities’ territories. BPCP staff follow strict hygiene and observation distance rules to prevent disease transmission. Following the Covid19 outbreak, from March 2020 data collection moved from chimpanzee follows to biomonitoring only. Regular follows were resumed in 2022. Permissions to collect data were provided by the National Forest Authority of Uganda, the Uganda Wildlife Authority, and the Ugandan National Council of Science and Technology, and all data collection adhered to national and international guidelines, including the American Society of Primatologists Principles for the Ethical Treatment of Non-Human Primates, and the Code of Best Practices for Field Primatology.

## Results

### Presence of ground and tree nests

Between August 2019 and July 2022, we recorded 891 night-nests within 310 nesting sites, number of nests per site ranged from 1 to 21 (Figure 1). We recorded 692 tree nests and 189 ground nests (21%). Ground nests occurred in 78 of the 310 sites (25% of nest sites) and co-occurred with tree nests in 31 of these sites (10% all nest sites; 40% of sites with a ground nest; up to 15 ground nests within a site). Ground nests were observed year-round (50% in drier season months Dec-Feb, Jun-Aug; 50% in wetter season months Mar-May, Sep-Nov).

Between August 2019 and February 2020 we recorded additional information on nest construction and local habitat for 138 nesting sites containing 323 night-nests (282 tree nests, 41 ground nests). Tree nests varied in height from 1-45m; and n=11 (4%) were at 2m or less from the ground. All ground nests that could be measured (n=39) contained detached interwoven branches, and most (n=34, 87%) incorporated bent branches as well (see Figure 2). A small number (n=2) incorporated terrestrial herbs into the construction. The majority of nest sites were located in medium-density forest (n=117) as opposed to open (n=17) or high-density (n=4); similarly, the majority of nest sites were located in mixed-secondary forest (n=121) rather than disturbed (n=8) or savannah (n=1; Unknown n=8).

**Figure 2:**
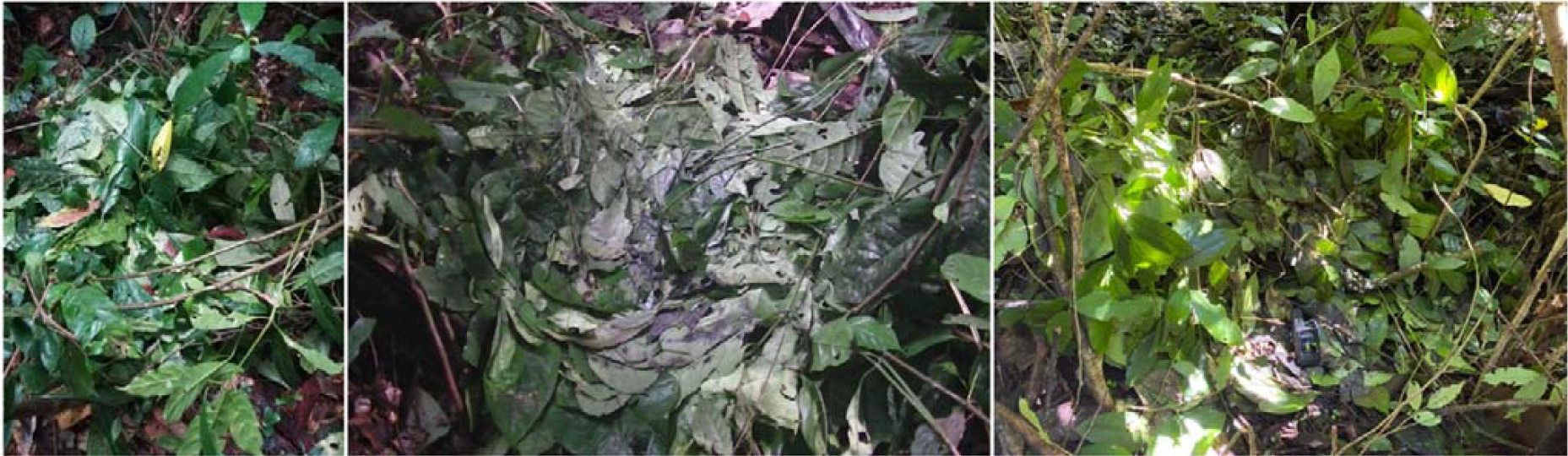
Three ground nests. Pictures of three ground nests at three separate nest sites. Note the inclusion of multiple bent and interwoven branches together with substantial quantities of smaller leafy material, distinguishing them from flimsier resting structures such as ‘day beds’ and ‘cushions’.

## Discussion

Bugoma chimpanzees regularly construct ground nests overnight and, in addition, construct tree nests at a height that would be vulnerable to human threat (<2m). In addition to their observed frequency, we find ground nests most often appeared as clusters suggesting that they were not built by a single individual or a single family. As these chimpanzees were not fully habituated at the time of data acquisition, direct observations of overnight ground nest building were not possible. The estimated number of independent individuals in the community (∼40) are a fraction of the total number of ground nests encountered, so it is impossible that each ground nest observed was built by a separate individual on only one occasion. Similarly, up to 15 ground nests were found in a single nest site, thus, it cannot be the case that these are the work of a single extremely prolific individual. We would suggest that the most likely explanation is the repeated building of ground nests by at least some individuals. If so, ground nesting in the Mwera community of chimpanzees meets the criteria for a ‘habitual’ behaviour (Whiten et al., 1999).

The transition to regularly sleeping on the ground has been suggested as an important driver of hominin behavioural-cognitive changes in our divergence from other apes (Coolidge & Wynn, 2006) and, thus, the factors that drive more regular occurrence of ground-sleeping in apes are of interest across diverse fields. It is possible that local ecological factors promote the use of ground nesting in the Bugoma communities. While systematic research is needed to confirm this, we believe that limitations in the number of suitable trees for tree nesting (c.f., McCarthy et al. 2016) is unlikely to be driving ground nesting, both because of the regular co-occurrence of tree and ground nests in the same nesting party; and because the secondary mixed-forest habitat in which the majority of nest sites are located is similar in structure to that of the neighbouring Budongo Forest Reserve, in which no overnight ground nests have been reported in over 40-years of cumulative observation across two communities.

Another explanation could be seasonality (Pruetz et al. 2018; Tagg et al., 2013), but Bugoma chimpanzees seem to nest on the ground year-round (see also Koops et al. 2012). Nesting on the ground may provide important differences in thermoregulation (Stewart, 2011b) or in pathogen avoidance (Stewart, 2011b; Videan, 2006; Lacroux et al., 2022), which are of substantial interest for future investigation in Bugoma; however, the frequent co-occurrence of ground and tree nests in the same nest site suggests that any effect here varies by individual characteristics. One possibility is that there are sex-specific differences: for example, ground nesting may be more prevalent in male chimpanzees, who already typically nest at lower heights (e.g., Brownlow et al., 2001). Chimpanzees also appear to select nesting tree species for ‘comfort’, taking into account features such as high leaf density (Lacroux et al., 2023), an aspect that could also vary by individual or sex.

Another ecological explanation might be that large numbers of individuals are unable to construct tree nests due to injury or illness. Again, we believe that this is unlikely. Our long-term data collection includes health monitoring of the chimpanzees and while there are outbreaks of respiratory infection in some individuals, these appear similar to those reported in other Uganda forest populations (Scully et al., 2018). A more likely pressure would be the impact of snare wires which regularly maim chimpanzees who trap their hands and feet in them – these injuries can include wasted and amputated limbs. However, once again, the neighbouring Budongo communities also have high numbers of snare-injured chimpanzees (Reynolds, 2005; Stokes & Byrne, 2006; Waller & Reynolds, 2001), including individuals missing hands and feet (including individuals where more than one limb is amputated), and do not report any use of ground nesting.

Ground nesting has sometimes been explained by scholars as resulting from an absence of predators or human activity (Pruetz et al. 2008; Hicks, 2010; McCarthy et al., 2012; Last & Muh, 2013). In Bugoma a number of features of the forest make ground nesting potentially risky. While there are no recent confirmed sightings of chimpanzees’ main nonhuman predator (leopard, *Panthera pardus*) in Bugoma, other large animals that could present potential risk were regularly directly or indirectly (through camera-traps) observed over the study period, including elephant (*Loxodonta cyclotis*), golden cat (*Caracal aurata*), python (*Python sebae*), and rhinoceros viper (*Bitis nasicornis*; Figure 3). Elephant presence has been previously described as impacting ground nesting at Lope (Tutin et al., 1995) and they are also present other sites where ground nesting has been observed (i.e., La Belgique, Tagg et al., 2013; Bili-Uele, Hicks 2010). Perhaps the most significant risk to chimpanzees in Bugoma is human activity. Hunting is prominent; as well as the use of snare wires designed to target duiker and pig species, hunters in Bugoma are regularly encountered with large packs of dogs in the chimpanzees’ range (Figure 4). The butchered remains of chimpanzees, apparently following human hunting, have been discovered by the project on two occasions, with additional unconfirmed reports (Bugoma Primate Conservation Project, unpublished data). Interestingly, while it remains counter-intuitive, Tagg and colleagues (2013) also found that chimpanzee ground nesting appeared to increase with human pressure.

**Figure 3:**
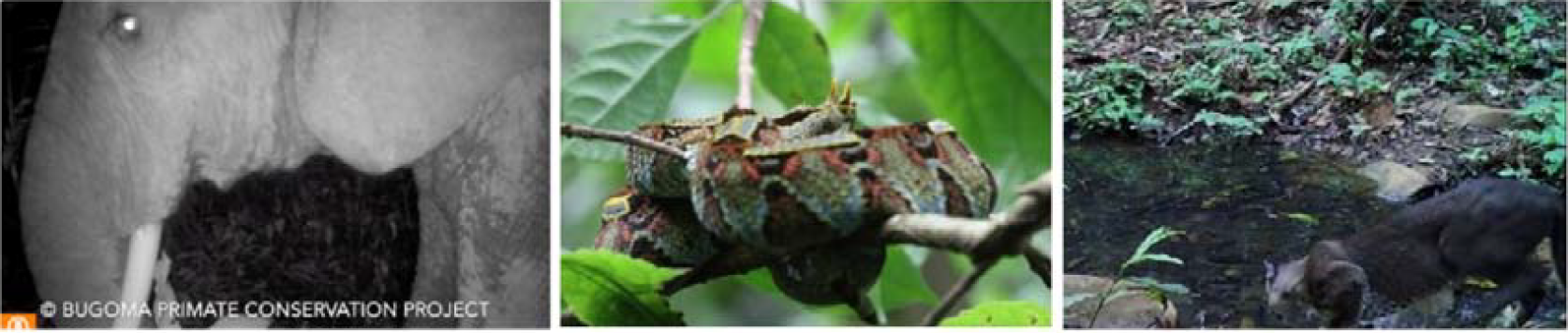
Large primarily terrestrial mammals and reptiles present in the Bugoma Central Forest Reserve that represent a potential threat to chimpanzees. Pictures captured directly or by camera trap within the home range of the Mwera community. From Left to right: African elephant (*Loxodonta cyclotis)*; Rhinoceros viper (*Bitis nasicornis)*; Golden cat (*Caracal aurata)*.

**Figure 4:**
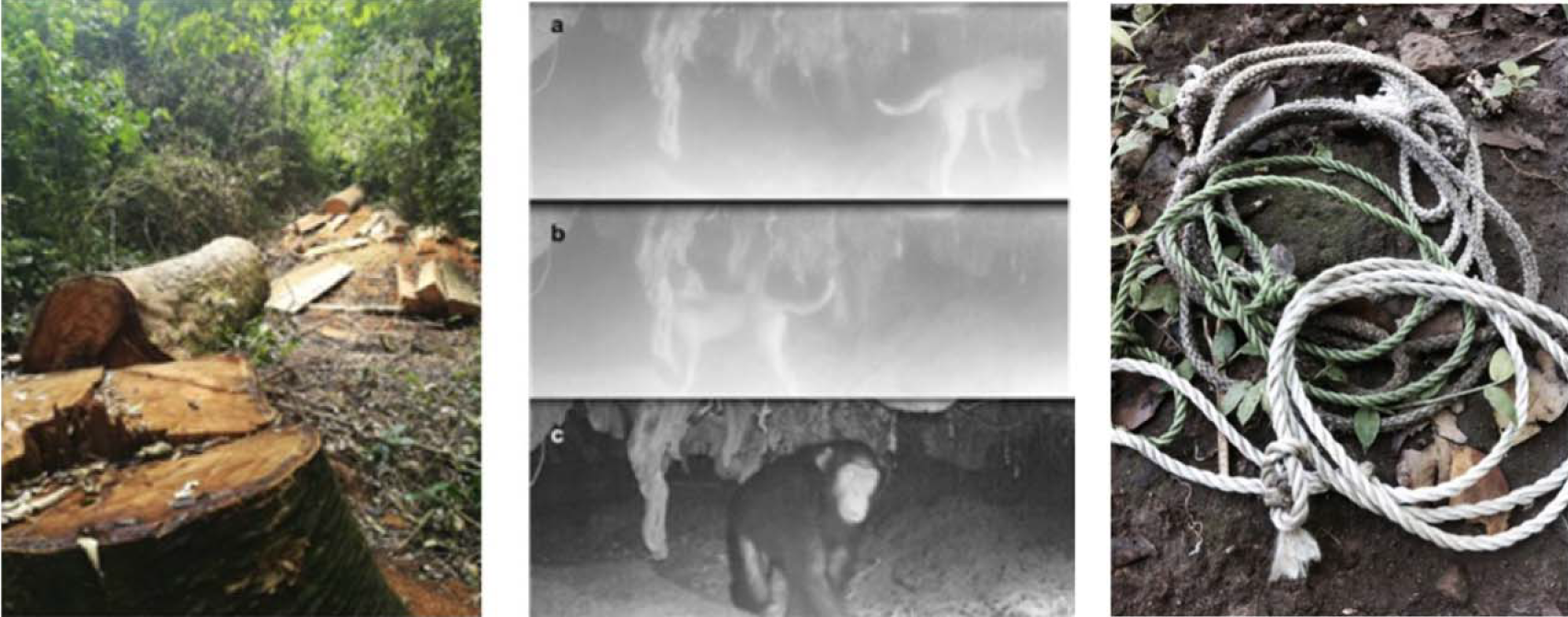
Human activities in the Bugoma Central Forest Reserve. Pictures captured directly or by camera trap in the home range of the Mwera community. Left to right: illegal timber felling; overlapping presence of hunting dogs in key resource locations for chimpanzees; use of wire and nylon snares for trapping small game.

While substantial further research is needed, we do not, as yet, have clear evidence for any ecological or physical drivers of ground nesting in Bugoma. Given the significant risk to life by, at least, human hunters, it is possible that there is a cultural component to the Bugoma chimpanzee ground nesting. If true, this would highlight that, as in other species (Laland & Williams, 1998; Franz & Matthews, 2010), some socially acquired chimpanzee behavioural variants may not have a positive effect on individual fitness. Behavioural variants linked to improving vital rates, such as survival, were suggested as being of particular conservation importance (Brakes et al., 2021), but establishing these connections is challenging (Carvalho et al., 2022) and association with individual fitness may vary with fluctuating local pressures, for example human hunting. Chimpanzee cultures have been shown to be inflexible, with resistance to the modification or acquisition of new behaviour even where it could be advantageous (Boesch & Boesch-Achermann, 2000; Gruber et al., 2011; but see Hobaiter et al., 2014), suggesting that they may be similarly resistant to its loss where it appears to be disadvantageous.

With increasing human pressure on the forest, it is unclear how the Bugoma chimpanzees will react with respect to their nest building behaviour. Closely monitoring ground nest building in Bugoma forest may provide a direct assessment of the impact of anthropogenic activities on wild semi-habituated chimpanzees. The importance of community and population specific behavioural variation from a conservation perspective has started to receive significant attention (Brakes et al., 2019; Brakes et al., 2021). The loss of a specific community involves the irreversible loss of a unique genetic heritage, as well the information and individual knowledge that any local culture is founded on. The wider Bugoma forest habitat is under considerable pressure, not only from illegal activities such as hunting and logging, but also from clear-felling for crop production, from local energy resource extraction, and from increased infrastructure and anthropogenic activities (McLennan et al., 2020; Plumptre et al., 2020). Non-human behavioural variation is sensitive to the impact of human anthropogenic activity on the transformation, fragmentation, and degradation of habitats (Gruber et al., 2019; Kalan et al., 2020). Widespread urgent calls to action throughout 2019 to 2022 for its protection by Ugandan and international organisations, as well as planned sustainable resource generation from eco-tourism projects may benefit from a clear and strong case for the unique behavioural repertoire of the chimpanzees in this area.

## Abbreviations

GPS: Global Positioning System
BPCP: Bugoma Primate Conservation Project
m: meters

## Acknowledgements

We thank the staff of the Bugoma Primate Conservation Project, and the Ugandan National Forestry Authority and the Jane Goodall Institute Switzerland for their support of the field site. Fieldwork for this research received funding from the European Union’s 8^th^ Framework Programme, Horizon 2020, under grant agreement no 802719; National Geographic (Grant GS-63895R-19), the Swiss National Science Foundation (Grant PCEFP1_186832). Funders had no role in the study design, writing, or decision to publish. We thank the editor and two reviewers for their constructive comments that improved the manuscript.

## Conflict of Interest Statement

The authors declare they have no conflict of interest.

## Notes

https://github.com/Wild-Minds/GroundNesting_Bugoma

### Competing Interest Statement

The authors have declared no competing interest.

### Summary of Updates

To accommodate reviewer feedback we have edited for clarity, and provided a more cautious assessment of the possible cultural underpinnings - providing a more balanced discussion of the ecological and social drivers that may shape behavioural variation in nest building in Bugoma.

## References

Anderson, J. R. (1998). Sleep, sleeping sites, and sleep-related activities: awakening to their significance. American Journal of Primatology, 46, 63–75.

Anderson J.R., Ang M.Y.L., Lock M.C., & Weiche, I. (2019). Nesting, sleeping, and nighttime behaviors in wild and captive great apes. Primates, 60, 321–332. 10.1007/s10329-019-00723-2

Baldwin, P.J. (1979). The natural history of the chimpanzee (Pan troglodytes verus) at Mt. Assrik, Senegal. (Doctoral dissertation, University of Stirling).

Boesch, C. (1995). Innovation in wild chimpanzees (Pan troglodytes). International Journal of Primatology, 16(1), 1–16.

Boesch, C., & Boesch-Achermann, H. (2000). The chimpanzees of the Taï forest: Behavioural ecology and evolution. Oxford: Oxford University Press.

Brakes, P., Dall, S. R., Aplin, L. M., Bearhop, S., Carroll, E. L., Ciucci, P., … Rutz, C. (2019). Animal cultures matter for conservation. Science, 363, 1032–1034.

Brakes, P., Carroll, E. L., Dall, S. R., Keith, S.A., McGregor, P. K., Mesnick, S. L., … Garland, E. C. (2021). A deepening understanding of animal culture suggests lessons for conservation. Proceedings of the Royal Society B: Biological Sciences, 288, rspb.2020.2718

Brownlow, A. R., Plumptre, A. J., Reynolds, V., & Ward, R. (2001). Sources of variation in the nesting behavior of chimpanzees (Pan troglodytes schweinfurthii) in the Budongo Forest, Uganda. American Journal of Primatology, 55(1), 49–55.

Brugiere, D., & Sakom, D. (2001). Population density and nesting behaviour of lowland gorillas (Gorilla gorilla gorilla) in the Ngotto forest, Central African Republic. Journal of Zoology, 255(2), 251–259.

Carvalho, S., Wessling, E. G., Abwe, E. E., AlmeidaLWarren, K., Arandjelovic, M., Boesch, C., … & Koops, K. (2022). Using nonhuman culture in conservation requires careful and concerted action. Conservation Letters, 15(2), e12860.

Coolidge, F. L., & Wynn, T. (2006). The effects of tree-to-ground sleep transition in the evolution of cognition in early Homo. Before Farming, 4, 1–18.

Estrada, A., Garber, P. A., Rylands, A. B., Roos, C., Fernandez-Duque, E., Di Fiore, A., … Li, B. (2017). Impending extinction crisis of the world’s primates: Why primates matter. Science Advances, 3(1), e1600946.

Franz, M., & Matthews, L. J. (1998) Social enhancement can create adaptive, arbitrary, and maladaptive cultural traditions. Proceedings of the Royal Society B: Biological Sciences doi:10.1098/rspb.2010.0705

Fruth, B. (1995). Nests and nest groups in wild bonobos (Pan paniscus): Ecological and behavioural correlates. Aachen: Verlag Shaker.

Fruth, B., & Hohmann, G. (1993). Ecological and behavioral aspects of nest building in wild bonobos (Pan paniscus). Ethology, 94(2), 113–126.

Fruth, B., & Hohmann, G. (1996). Comparative analyses of nest building behavior in Bonobos and Chimpanzees. In: Wrangham, R.W, W.C. McGrew, F.B.M. de Wall & P.G. Heltne (eds). Chimpanzee Cultures. Harvard University Press: xxii pp 424.

Fruth, B., Tagg, N., & Stewart, F. (2018). Sleep and nesting behavior in primates: A review. American Journal of Physical Anthropology, 166(3), 499–509.

Furuichi, T., & Hashimoto, C. (2000). Ground beds of chimpanzees in the Kalinzu Forest, Uganda. Pan Africa News, 7, 26–28.

Galef, B. G., & Whiten, A. (2017). The comparative psychology of social learning. In J. Call, G. M. Burghardt, I. M. Pepperberg, C. T. Snowdon, T. Zentall (Eds.), APA handbook of comparative psychology: perception, learning, and cognition. pp. 411–439. APA.

Goodall, J. (1962). Nest building behavior in the free ranging chimpanzees. Annals of the New York Academy of Sciences, 102 (2): 455–467.

Guislain, P., & Dupain, J. (2005). Scientific report: determinants of habitat use by sympatric chimpanzee and gorilla populations at the periphery of the Dja Faunal Reserve, Cameroon. Report for the L.S.B. Leakey Foundation, PGS Cameroon.

Gruber, T., Muller, M. N., Reynolds, V., Wrangham, R. W., & Zuberbühler, K. (2011). Community-specific evaluation of tool affordances in wild chimpanzees. Scientific Reports, 1, 1–7.

Gruber, T., Luncz, L. V., Mörchen, J., Schuppli, C., Kendal R. L., & Hockings, K. J. (2019). Cultural change in animals: a flexible behavioural adaptation to human disturbance. Palgrave Communications 5, 1–9. doi: 10.1057/s41599-019-0271-4

Hernandez-Aguilar, R. A. (2009). Chimpanzee nest distribution and site reuse in a dry habitat: implications for early hominin ranging. Journal of Human Evolution, 57(4), 350–364.

Hicks, T. C. (2010). A chimpanzee mega-culture? Exploring behavioral continuity in Pan troglodytes schweinfurthii across northern DR Congo. African Primates, 7(1), 1–18.

Hicks, T. C., Kühl, H. S., Boesch, C., Dieguez, P., Ayimisin, A. E., Fernandez, R. M., … Roessingh, P. (2019). Bili-Uéré: A chimpanzee behavioural realm in Northern Democratic Republic of Congo. Folia Primatologica, 90, 3–64.

Hobaiter, C., Poisot, T., Zuberbühler, K., Hoppitt, W., & Gruber, T. (2014). Social network analysis shows direct evidence for social transmission of tool use in wild chimpanzees. PloS Biology, 12(2), e1001960.

Izawa, K., & Itani, J. (1966). Chimpanzees in Kasakati Basin, Tanganyika (1) Ecological study in the rainy season 1963–1964. Kyoto University African Studies 1, 73–156.

Kalan, A. K., Kulik, L., Arandjelovic, M., Boesch, C., Haas, F., Dieguez, P., … Kühl, H. S. (2020). Environmental variability supports chimpanzee behavioural diversity. Nature Communications, 11.1, 1–10. doi: 10.1038/s41467-020-18176-3

Koops, K., McGrew, W. C., de Vries, H., & Matsuzawa, T. (2012). Nest-building by chimpanzees (Pan troglodytes verus) at Seringbara, Nimba Mountains: antipredation, thermoregulation, and antivector hypotheses. International Journal of Primatology, 33, 356–380.

Koops, K., Humle, T., Sterck, E. H. M., & Matsuzawa, T. (2007). Ground-nesting by the chimpanzees of the Nimba Mountains, Guinea: environmentally or socially determined? American Journal of Primatology, 69, 407–419.

Kortlandt, A. (1992). On Chimpanzee Dormitories and Early Hominid Home Sites. Current Anthropology, 33, 399–401.

Lacroux, C., Pouydebat, E., Rossignol, M., Durand, S., Aleeje, A., Asalu, E., … & Krief, S. (2022). Repellent activity against Anopheles gambiae of the leaves of nesting trees in the Sebitoli chimpanzee community of Kibale National Park, Uganda. Malaria journal, 21(1), 1–11.

Lacroux, C., Krief, S., Douady, S., Cornette, R., Durand, S., Aleeje, A., … & Pouydebat, E. (2023). Chimpanzees select comfortable nesting tree species. Scientific Reports, 13(1), 16943.

Laland, K. N., & Williams, K. (1998). Social transmission of maladaptive information in the guppy. Behavioural Ecology, 9, 493–499.

Last, C., & Muh, B. (2013). Effects of human presence on chimpanzee nest location in the Lebialem-Mone Forest landscape, Southwest Region, Cameroon. Folia Primatologica, 84, 51–63

Lethmate, J. (1977). Nest-building behaviour of a young Orang-utan raised in isolation. Primates, 18(3), 545–554.

Matsuzawa, T., & Yamakoshi, G. (1996). Comparison of chimpanzee material culture between Bossou and Nimba, West Africa. In A. E. Russon, K. A. Bard, & S. T. Parker (Eds.), Reaching into thought: The minds of the great apes (pp. 211–232). Cambridge University Press.

McCarthy, M. S., Lester, J. D., & Stanford, C. B. (2017). Chimpanzees (Pan troglodytes) flexibly use introduced species for nesting and bark feeding in a human-dominated habitat. International Journal of Primatology, 38(2), 321–337

McGrew, W. C. (2004). The cultured chimpanzee: Reflections on cultural primatology. Cambridge: Cambridge University Press.

McGrew, W. C. (2010). Chimpanzee technology. Science, 328(5978), 579–580.

McLennan, M. R., Hintz, B., Kiiza, B. V., Rohen, J., Lorenti, G. A., & Hockings, K. J. (2020). Surviving at the extreme: chimpanzee ranging is not restricted in a deforested human-dominated landscape in Uganda. African Journal of Ecology, doi:10.1111/aje.12803

Mfossa, D. M., Gazagne, E., Gray, R. J., Ketchen, M. E., Abwe, E. E., Beudels-Jamar, R. C., … & Brotcorne, F. (2022). Montane Grassland Resources Drive Gorilla (Gorilla Gorilla) Nesting Behaviours in the Ebo Forest, Littoral Region, Cameroon. ResearchSquare DOI: 10.21203/rs.3.rs-2082431/v1

Morgan, D., Sanz, C., Onononga, J. R., & Strindberg, S. (2016). Factors Influencing the Survival of Sympatric Gorilla (Gorilla gorilla gorilla) and Chimpanzee (Pan troglodytes troglodytes) Nests. International Journal of Primatology, 37(6), 718–737. 10.1007/s10764-016-9934-9

Plumptre, A. J., & Reynolds, V. (1997). Nesting behavior of chimpanzees: implications for censuses. International journal of primatology, 18(4), 475–485

Plumptre, A. J., & Cox, D. (2006). Counting primates for conservation: primate surveys in Uganda. Primates, 47, 65–73.

Plumptre, A. J., Ayebare, S., Kujirakwinja, D., & Segan, D. (2020). Conservation planning for Africa’s Albertine Rift: conserving a biodiverse region in the face of multiple threats. Oryx, 55, 302–310.

Prasetyo, D., Ancrenaz, M., Morrogh-Bernard, H.C., Utami Atmoko, S.S., Wich, S.A., van Schaik, C.P. (2009). Nest building in orangutans. In: Wich, S.A., Utami Atmoko, S.S., Mitra Setia, T., van Schaik, C.P. Orangutans: geographic variation in behavioral ecology and conservation. New York, US: Oxford University Press, 269–277.

Pruetz, J., Fulton, S. J., Marchant, L. F., McGrew, W. C., Schiel, M., & Waller, M. (2008). Arboreal nesting as anti-predator adaptation by savanna chimpanzees (Pan troglodytes verus) in southeastern Senegal. American Journal of Primatology, 70, 393–401.

Reynolds, V. (2005). The chimpanzees of the Budongo forest: Ecology, behaviour and conservation. Oxford University Press, Oxford.

Romani, T., Mundry, R., Mayanja Shaban, G., Konarzewski, M., Namaganda, M., Hobaiter, C., Gruber, T., & Hicks, T. C. (2023). Decay rates of arboreal and terrestrial nests of eastern chimpanzees (Pan troglodytes sweinfurthii) in the Bugoma Central Forest Reserve, Uganda: implications for population size estimates. American Journal Primatology.

Samson, D. R., & Hunt, K. D. (2012). A thermodynamic comparison of arboreal and terrestrial sleeping sites for dry-habitat chimpanzees (Pan troglodytes schweinfurthii) at the Toro-Semliki Wildlife Reserve, Uganda. American Journal of Primatology, 4, 811–818.

Samson, D. R., & Hunt, K. D. (2014). Chimpanzees preferentially select sleeping platform construction tree species with biomechanical properties that yield stable, firm, but compliant nests. PLoS One, 9, e95361.

Scully, E. J., Basnet, S., Wrangham, R. W., Muller, M. N., Otali, E., Hyeroba, D., … Goldberg, T. L. (2018). Lethal respiratory disease associated with Human Rhinovirus C in wild chimpanzees, Uganda, 2013. Emerging Infectious Diseases, 24, 267–274.

Schaller, G. E. (1963). The Mountain gorilla: Ecology and behavior. Oxford, England: University of Chicago Press.

Stewart, F. A. (2011). The evolution of shelter: ecology and ethology of chimpanzee nest building (Doctoral dissertation, University of Cambridge).

Stewart, F. A. (2011b). Why sleep in a nest? Empirical testing of the function of simple shelters made by wild chimpanzees. American Journal of Biological Anthropology, 146(2), 313–318.

Stewart, F. A., & Pruetz, J. D. (2013). Do chimpanzee nests serve and anti-predatory function? American Journal of Primatology, 75, 593–604.

Stewart, F. A., Piel, A. K., Azkarate, J. C., & Pruetz, J. D. (2018). Savanna chimpanzees adjust sleeping nest architecture in response to local weather conditions. American Journal of Physical Anthropology, 166(3), 549–562.

Stokes, E. J., & Byrne, R. W. (2006). Effect of Snare Injuries on the Fig-Feeding Behavior of Chimpanzees of the Budongo Forest, Uganda. In Primates of western Uganda (pp. 281–297). Springer, New York, NY.

Sylla, S. F., Ndiaye, P., Lindshield, S. M., Bogart, A. L., & Pruetz, J. D. (2022). The western chimpanzee (pan troglodytes verus) in the antenna zone (niokolo koba national park, senegal): nesting ecology and sympatrics with other mammals. Applied ecology and environmental research, 20(3), 2663–2681.

Suzuki, A. (1969). An ecological study of chimpanzees in a savanna woodland. Primates, 10(2), 103–148.

Tagg, N., Willie, J., Petre, C.A., Haggis, O. (2013). Ground night nesting in chimapnzees: new insights from central chimpanzees (Pan troglodytes troglodytes) in south-east Cameroon. Folia Primatologica, 84(6), 362–383.

Tutin CEG, Fernandez M (1984). Nationwide census of gorilla (Gorilla g. gorilla) and chimpanzee (Pan t. troglodytes) populations in Gabon. American Journal of Primatology, 6, 313–336.

Tutin, C. E., Ham, R., & Wrogemann, D. (1995). Tool-use by chimpanzees (Pan t. troglodytes) in the Lopé Reserve, Gabon. Primates, 36(2), 181–192.

van Lawick-Goodall, J. (1968). The behaviour of free-living chimpanzees in the Gombe Stream Reserve. Animal Behaviour Monographs, 1, 161–311.

van Casteren, A., Sellers, W. I., Thorpe, S. K., Coward, S., Crompton, R.H., Myatt, J. P., & Ennos, A. R. (2012). Nest-building orangutans demonstrate engineering know-how to produce safe, comfortable beds. Proceedings of the National Academy of Sciences, 109, 6873–6877.

Videan EN (2006) Bed-building in captive chimpanzees (Pan troglodytes): the importance of early rearing. American Journal of Primatology, 68, 745–751.

Waller, J. C., & Reynolds, V. (2001). Limb injuries resulting from snares and traps in chimpanzees (Pan troglodytes schweinfurthii) of the Budongo Forest, Uganda. Primates, 42(2), 135–139.

Wessling, E. G., & Surbeck, M. (2021). Failure to account for behavior variability significantly compromises accuracy in indirect population monitoring. Animal Conservation.

Whiten, A., Goodall, J., McGrew, W. C., Nishida, T., Reynolds, V., Sugiyama, Y., … Boesch, C. (1999). Culture in chimpanzees. Nature, 399, 682–685.

Wilson, M. L., Boesch, C., Fruth, B., Furuichi, T., Gilby, I. C., Hashimoto, C., … & Wrangham, R. W. (2014). Lethal aggression in Pan is better explained by adaptive strategies than human impacts. Nature, 513(7518), 414–417.

